# “Rapid Eye Movement sleep deprivation of rat generates ROS in the hepatocytes and makes them more susceptible to oxidative stress”

**DOI:** 10.1101/375683

**Authors:** Atul Pandey, Santosh K Kar

## Abstract

**Background:** Rapid Eye Movement sleep deprivation (REMSD) of rats causes inflammation of the liver and apoptotic cell death of neurons and hepatocytes. Studies also suggest that REMSD are involved with muscle injury, cardiac injury and neurodegerative diseases.

**Objective and methods:** The aim of this research was to determine whether REMSD of rats would generate reactive oxygen species (ROS) and create oxidative stress in the hepatocytes. We selectively deprived the rats from REM sleep using the standard flower pot method.

**Results:** We observed that when rats were subjected to REMSD, the levels of ROS in the hepatocytes increased with the increase in the number of days of REMSD by ∼265%, but it returned towards normal levels after recovery sleep for 5 days (∼36%) compared to controls. Nitric oxide synthase (iNOS) gene and protein was found elevated in hepatocytes in response to REM sleep loss as confirmed by real time PCR and western blot analysis compared to controls. The level of nitric oxide (NO) also increased by ∼ 675% in the hepatocytes of REMSD rats as compared to that of control group of animals.

**Discussion:** We have analyzed the oxidative stress generated and potentiation of hepatocytes against oxidative stress in response to REMSD. Since, REM sleep is known to play an important role for survival of most animals and has important role in maintenance of body physiology. Hence, our findings that loss of REM sleep in hepatocytes of rats can affect the ROS levels and induce iNOS & NO circulation, while making them more susceptible to oxidative stress assumes significance.

**Highlights of the study:**

- We observed elevated levels of ROS in the hepatocytes of REM sleep deprived rats.
- The hepatocytes of REMSD group of rats were found more susceptible to oxidative stress than that of control groups.
- We found increased expression of iNOS gene and nitric oxide synthase protein in the hepatocytes of REMSD rats.
- We observed that nitric oxide levels in the hepatocytes of REM sleep deprived rats increased positively with days of REMSD, but returned to its normal levels after 5 days of recovery sleep.

## Introduction

Sleep is a natural physiological process which is needed for the survival of most living beings studied so far[1]. Sleep is an enigma and is known to be involved in healing processes and repair of our heart and blood vessels and its deficiency is linked to increased risk of heart and kidney diseases, high blood pressure, diabetes and stroke[2–5]. In mammals, sleep is broadly categorized as two types, namely non-rapid eye movement sleep (NREM) and rapid eye movement sleep (REM). REM sleep is present in all mammals and birds and is associated with maintenance of certain essential body functions and ability to survive under different ecological conditions. Besides, it helps in memory consolidation [6,7], brain maturation, spatial memory acquisition [8] and maintenance of body physiology[9]. Prolonged loss of REM sleep can alter blood brain barrier functions[10] and can be fatal [11,12]. In terms of energy metabolism, there is no difference between wakefulness and when the animal is in REM sleep, which represents approximately 1/3^rd^ of total sleep [13], But extended sleep deprivation experiments in animals have shown that during sleep there is decreased cerebral glucose utilization [14].

Most of studies on REMSD have focused on analyzing the biochemical changes that takes place in the brain and neurons and very little attention has been given to study other organs. Recent studies have shown that REMSD can cause apoptosis of neurons in the brain and can cause muscle and cardiac injuries [11-13]. But, whether total or partial sleep deprivation can cause damage to other organs of the body has remained unanswered. Reactive oxygen species (ROS) are generated when metabolic processes takes place in the tissues and body has a natural system to maintain the balance using anti-oxidants present in the system or available from dietary sources. The production of excessive ROS due to uncontrolled metabolic processes under stressful conditions can lead to a state where maintenance and recovery to normal physiological levels becomes difficult. This can ultimately lead to cell death and tissue disintegration. Studies performed for testing the relationship between sleep and restoration of normal functioning of the body has primarily focused on total sleep deprivation. There has been several studies giving contradictory reports, some indicating that ROS are generated due to sleep deprivation and are responsible for alteration of body physiology while others show no evidence of generation of ROS. One study supporting the role of ROS has reported that lipid peroxidation levels increased in the hippocampus of rodents due to increased ROS levels generated due to sleep deprivation[16]. In another study the antioxidant level were found decreased in peripheral tissues of animals that were deprived of sleep for 5 and 10 days[17], but it was restored after rebound recovery sleep. There are reports of increased superoxide dismutase (SOD) activity in rats which were subjected to total sleep loss (3-14days) and then allowed to recover in comparison to control groups[18]. More studies suggests that sleep deprivation create stress like conditions coupled with oxidative stress leading to decreased levels of glutathione in the whole brain [19] and reduced SOD activity in the hippocampus and brain stem[20].

Interestingly, there exists contradicting report giving no evidence of oxidative damage either in the brain or peripheral tissues such as liver after either short term (8hrs) or long term (3-14 days) sleep deprivation and no increase in antioxidant enzymes like SOD, catalase, glutathione peroxidase or malondialdehyde activity was reported [21]. It has been reported that prolonged wakefulness activates an adaptive stress pathway termed as the unfolded protein response, which momentarily guards against the detrimental consequences of ROS [25, 26]. A recent report has shown a sharp increase in lipid peroxidation in the hepatocytes of sleep deprived rats [23]. Liver, being the metabolic hub, contributes significantly to the maintenance of body physiology [24]. Since hepatocytes are known to be involved in metabolism, detoxifying endo- and xenobiotics and synthesis of many proteins like albumin it will be interesting how REMSD will affect them.[25–27]. Hence, we required to know whether REMSD of rats would generate ROS in the hepatocytes of the liver and how the hepatocytes would respond to it.

Our previous study had shown that REMSD induced acute phase response in the liver, increased the synthesis of pro-inflammatory cytokines like Interleukin (IL)1β, IL-6 and IL-12 and increased the levels of liver toxicity marker enzymes, alanine transaminase and aspartic transaminase circulating in the blood [28]. In the present study, we show that REMSD of rats affects the level of ROS in the hepatocytes, increases the expression of inducible nitric oxide synthase gene and the levels of the corresponding protein and interestingly makes the hepatocytes more susceptible to oxidative stress. We further observed the increased production of nitric oxide by the hepatocytes in response to REMSD.

## Materials and Methods

Male wistar rats weighing between 220-260 gm were used in this study. Animals were housed in the institutional animal house facility with a 12:12 hr L: D cycle (lights on at 7.00 am). Food and water were provided *ad libtium.* All experiments were conducted as per the protocol approved by the University’s Institutional Animal Ethics Committee.

### Methods used for REM sleep deprivation and recovery

Animals were REM sleep deprived by the most widely used flower-pot method [29,30]. In this method, animals were kept on a small raised platform (6.5 cm diameter) surrounded by water compared to large platform control (LPC) rats which were maintained on a platform of 12.5 cm diameter. Cage control (CC) rats were maintained in cages under laboratory condition. Although the animals could sit, crouch and have non-REM sleep on the small platform (6.5 cm diameter), but due to the postural muscle atonia during REM sleep, they were unable to maintain their extended relaxed body posture on the small platform and tended to fall into the surrounding water. As a result they wake up at the onset or prior to the appearance of REM sleep and thus deprived of it. After stipulated days of deprivation the recovery groups of rats were allowed to sleep uninterrupted in individual cages with sufficient supply of food and water. The rats were sacrificed on different days (4 & 9 days) of REMSD and after 5 days of REMSD recovery and hepatocytes were isolated for further analysis.

### Hepatocytes preparation

Hepatocytes were isolated from the liver of male wistar rats from CC, LPC and REM sleep deprived groups by method used by Liu et.al;2002 [31]. Briefly, the abdomens of the rats were opened through a midline incision and liver was taken out. The portal cannula was then placed on suitable platform and liver was perfused with 0.02% EDTA solution at 37 degree C, at a flow rate of 30 ml per minutes for 15 minutes in a culture dish. Subsequently, the collagenase solution (37° C) was recirculated through the liver at the same flow rate for 15 minutes. After perfusion, liver capsules were disrupted and digested liver parenchyma was suspended in the ice-cold Hank’s balance salt solution. The resulting cell suspension was washed by centrifuging at 500rpm for 5 min 2-3 times and further centrifuged over 30% percoll at 100g for 5 min to obtain pure hepatocytes. Viability of the cells as measured by trypan blue exclusion was ≥ 95%.

### ROS measurement and optimization of dose for creating oxidative stress

ROS level in the hepatocytes obtained from control as well as REMSD group of rats was measured after labeling with 2’,7’-dichlorodihydrofluorescein diacetate (DCF-DA), which is a cell-permeable dye that becomes fluorescent upon reaction with ROS such as hydroxyl radical, hydrogen peroxide, or peroxynitrite. Briefly, hepatocytes from different groups of rats were treated with 10 µMDCF-DA for 30 min followed by either ROS measurement or treatment with different doses of H_2_O_2_ (50-500µM), Xanthine/Xanthine oxidase (X/XO; 10-100mU/ml) for ROS induction and measurement depending upon experimental design. Whenever, specific ROS inhibitors such as catalase (100-500U/ml) and oxypurinol (10-50mU/ml) were used the hepatocytes were treated with them first followed by DCF-DA treatment. Then the hepatocytes were incubated with H2O2 and X/XO to induce oxidative stress condition following the protocol described for mouse osteoblasts and Human MG63 cells[32]. The cells were harvested at the indicated time points after incubation, washed three times with phosphate-buffered saline, and then immediately analyzed by Flow cytometer with a488-nm excitation beam. The signals were obtained using a 530-nm band pass filter (FL-1 channel) for DCF. Each determination is based on the mean fluorescence intensity of 5,000 cells.

### Assay for hepatocellular NO production

Hepatocytes were seeded in24-well, flat-bottom plates (Falcon) at 1 × 10^5^ cells/well and incubated at 37°C in humidified 5%CO_2_ conditions with and without Lipopolysaccharides. NO was measured indirectly by determination of the concentration of nitrite. Briefly, 100-μl aliquots were removed from hepatocytes cultures and incubated with an equal volume of Griess reagent (1% sulfanilamide, 0.1% *N*-1-naphthylethylenediamide dihydrochloride in 2.5% phosphoric acid) for 10 min at 37°C. Absolute values were determined using sodium nitrite as standard. Absorbance was read at 550 nm on a microtiter plate reader.

### Isolation of RNA and TaqMan Real-time PCR

RNA was isolated from hepatocytes harvested from CC, LPC and REM sleep deprived groups and stored in RNA later (Sigma-Aldrich, Cat. R0901) using RNA purification kit (RNeasy Mini Kit, Qiagen, Germany). RNA concentrations and integrity was assessed by Nanodrop and Agilent 2100 Bioanalyzer (Agilent Technologies, Massy, France). Total RNA isolated from the hepatocytes were converted to cDNA using the reverse transcription PCR kit (Applied Biosystems, United States). GADPH served as a housekeeping gene while cage control group served as calibrator Probes (Reporter dye FAM labeled on 5’ end and quencher VIC labeled on 3’ end), PCR master mix, and PCR plate/tubes were obtained from Applied Biosystems and the manufacturer’s instruction were followed. The catalogue numbers of gene probes were glyceraldehyde 3 phosphate dehydrogenase (GAPDH, Rn01749022_g1), iNOS (Rn02132634_s1) and master mix (Rn99999916-g1).

### Western Blot Analysis

Western blotting was performed as previously described [33]. Primary antibody used were in 1/1000 dilutions (Santacruz Biotechnology, Inc, USA), whereas secondary antibody which is horseradish peroxidase-conjugated were used at 1/5000 dilutions (Santacruz Biotechnology, Inc, USA). After transfer of proteins membranes were developed using enhanced chemiluminescence (ECL) reagent (Promega) and photographed using software (Photo and Imaging 2.0; Hewlett-Packard). The analysis of the images was performed with imaging software (Photoshop 8.0; Adobe).

### Statistical Analysis

We used SPSS (Version 2.7.2) for one-way ANOVA and Tukey’s HSD posthoc test for measuring out the effect across treatment groups. The p values ≥ 0.05 were considered significant.

## RESULTS

### Optimization of dose for induction of ROS using H_2_O_2_ and X/XO and it’s suppression by catalase or oxypurinol

We treated normal hepatocytes with different amount of H_2_O_2_ & X/XO in order to generate superoxide anion. Intracellular ROS level increased in a dose dependent manner with treatments of H_2_O_2_ (50–500 µM, ANOVA, F=354.91, df=4, p<0.001) and X/XO (10–100 mU/ml, ANOVA, F=374.83, df=4, p<0.001) for 60 minutes (Fig. 1a &b). We selected the dose of 200µM for H_2_O_2_ and 50 mU/ml for X/XO for the treatments and induction of oxidative stress based on the dose optimization experiments. (Fig.1a &b). When we treated the normal hepatocytes using 200µM-H_2_O_2_and 50 mU/ml-X/XO for different time points ranging from 20-120 minutes, we observed that 60 minutes is quite effective and sufficient for induction of superoxide anions (Fig 1c, ANOVA, F=410.21, df=4, p<0.001& Fig.1d, ANOVA, F=864.77, df=4, p<0.001). Further, to establish the effective concentrations of the scavengers of the ROS, we first treated the normal hepatocytes with respective doses of catalase (100-500U/ml) or oxypurinol (10-50µM) followed by 200µM-H_2_O_2_ and 50 mU/ml-X/XO treatments. The doses of catalase (500U/ml, Fig 1e,f ANOVA, F=135.71,df=3,p<0.001) and oxypurinol (50µM, Fig.1f, ANOVA, F=128.36,df=3,p<0.001) were found to be quite effective against treatments of H_2_O_2_ (200µM) and X/XO (50 mU/ml) for 60 minutes and were used for further experiments. These results indicate that we selectively succeeded in inducing the ROS levels after H_2_O_2_ (200µM) and X/XO (50 mU/ml) treatments and catalase (500U/ml) or oxypurinol (50µM) were able to quench the induced levels of ROS.

**Figure 1:**
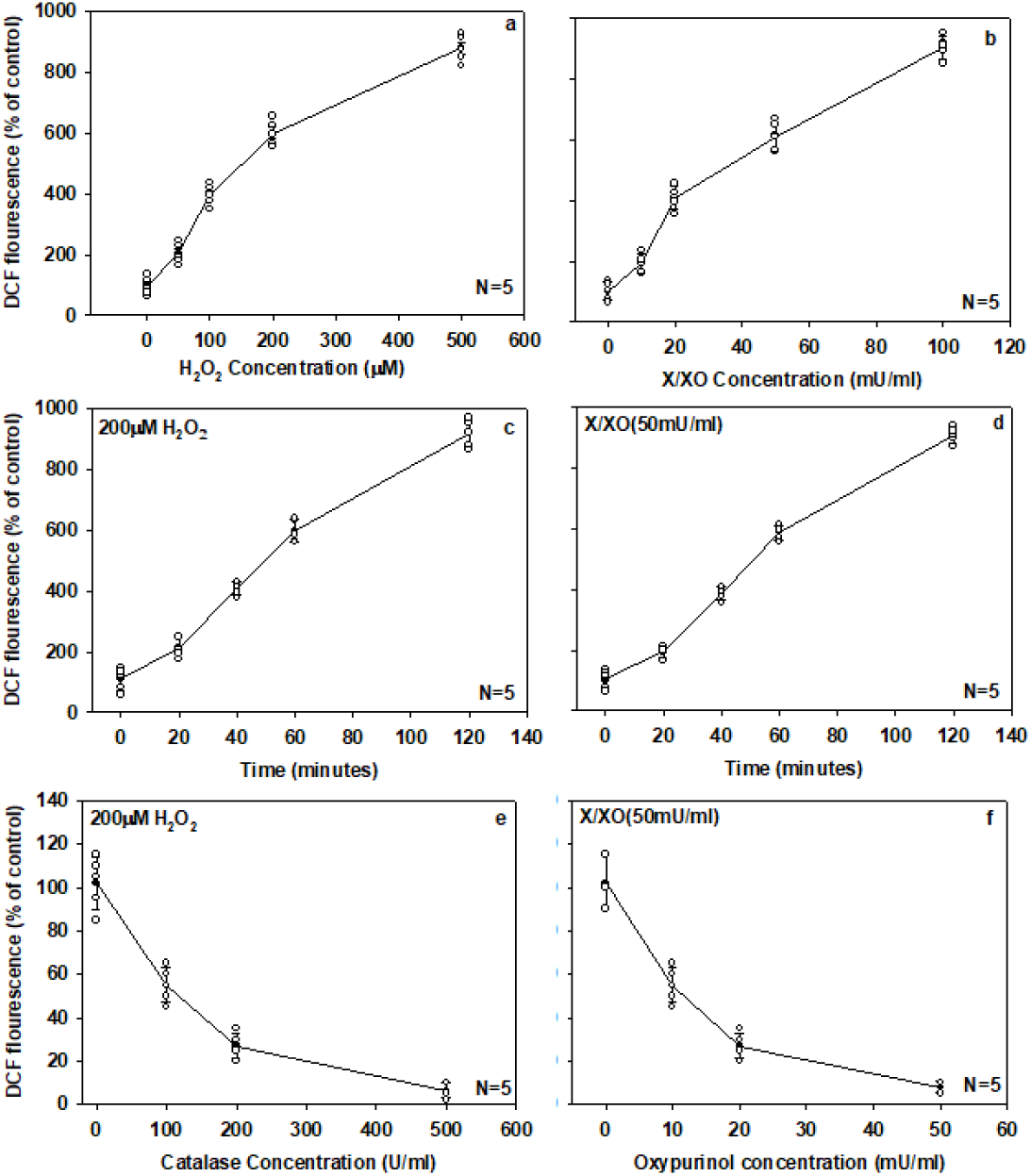
Optimization of ROS level induction and suppression in normal rat hepatocytes by H_2_O_2_, X/XO, catalase and oxypurinol. Hepatocytes, isolated from rat were loaded with 10 µM 2, 7-DCF-DA for 30 min and then treated with (a), 50–500 µM H_2_O_2_ for 1 h; (b), 10–100 µM xanthine (X) and 10–100 milliunits/ml Xanthine oxidase (XO) for 1 h; (c), 200 µM H_2_O_2_ for 20–120 min; (d), 50 µM-X and 50 milliunits/ml-XO for 20–120 min; (e), the hepatocytes were treated with different amount of catalase (100-500U/ml) after 200µM H_2_O_2_ induction for an hour and (f), scavenging of ROS production induced by 50mU X/XO for an hour using oxypurinol (10-50 mU/ml). ROS levels were then determined by FAC calibur as described under “Experimental Procedures.” Each point represents the mean ±S.E. of eight determinations from five (N=5) different cell samples. Each determination is the mean DCF fluorescence intensity of 5,000 cells.

### ROS production in hepatocytes by REM sleep deprivation

ROS production increased significantly with increase in the magnitude of REM sleep loss. We observed 111.15±12.5% increase in production of ROS in hepatocytes on day 4 which increased to 265.22±10.1% on day 9 compared to CC group treatment. The LPC group of rats showed comparable levels of ROS as in CC group of rats ruling out that changing from cage to large platform had any effect. Sleep recovery of 5 days showed decreased levels of ROS (36.36±5.1%) in comparison to CC group indicating the redressal effects (Fig. 2a, ANOVA F=216.92,df=27,p<0.001). At the same time, when we exposed the hepatocytes collected from different treatment group of rats with ROS quenchers (500U/ml catalase or 50µM oxypurinol) which are known to reduce the oxidative stresses, we got reduced levels (almost comparable to controls) of ROS in hepatocytes of 4 and 9 days REM sleep deprived rats (Fig.2a). This showed that ROS were generated in the hepatocytes due to REMSD and it got quenched by catalase or oxypurinol treatment.

**Figure 2:**
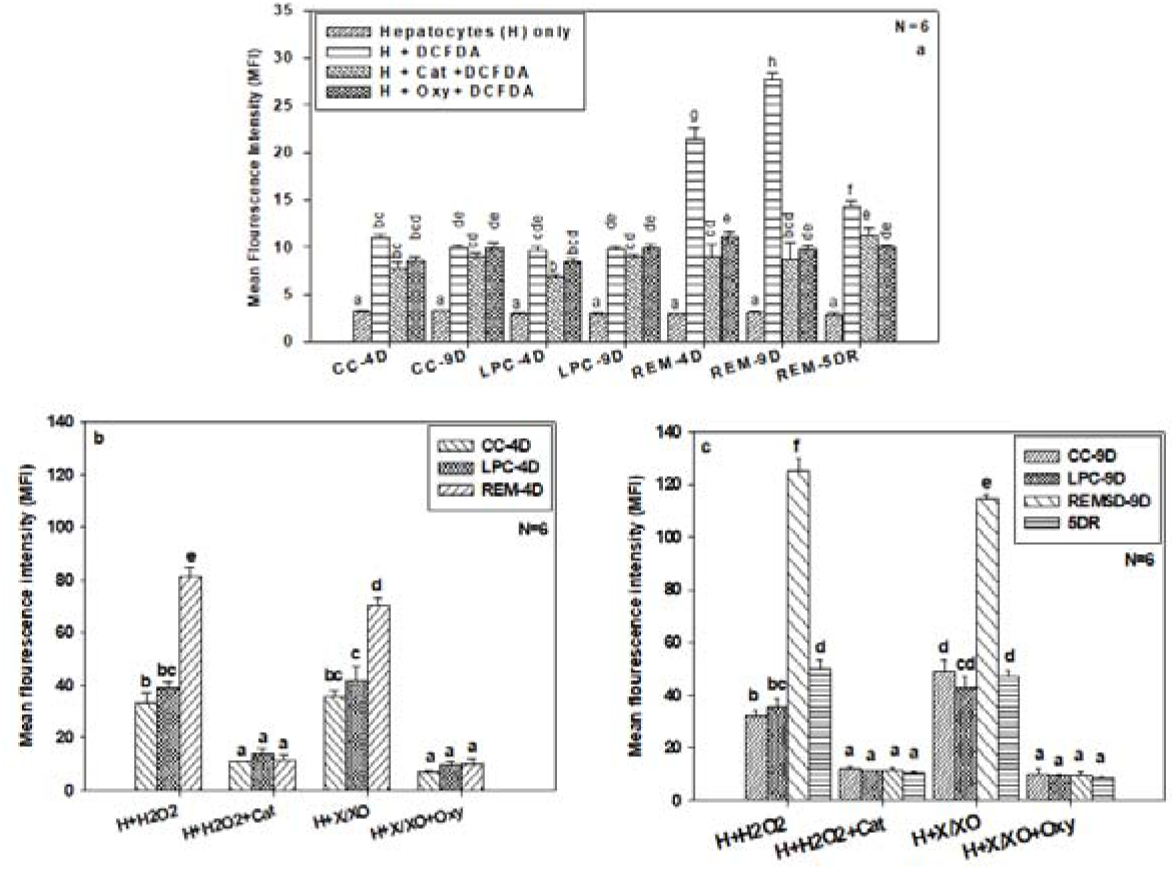
Effect of REM sleep deprivation on ROS production in hepatocytes. Measurement of ROS in hepatocytes from cage control, large platform control, REM sleep deprived group and recovery group of rats. (a), Measurement of ROS in different control groups and REM sleep deprived group of rats treated with scavengers like catalase and oxypurinol after 4 day, 9 days of REM sleep deprivation and 5day of sleep recovery. Differential productions of ROS in progression with REM sleep deprivation compared with control and simultaneously treated with catalase and oxypurinol after 4day of sleep deprivation (b), and 9 day of REMSD and 5day of sleep recovery after 9days of REMSD (c), similarly treated with catalase and oxypurinol. Each vertical bar represents the mean of 6 individual samples with standard error. Treatments that don’t share a letter, are statistically different in Tukey post-hoc analysis followed by one way ANOVA across treatment groups. [CC=Cage control, LPC=Large platform control, REM= Rapid eye movement sleep deprivation. H=Hepatocytes, DCFDA= dichlorodihydrofluorescein diacetate, Cat= catalase (200U/ml), Oxy=Oxypurinol (50µM), X/XO=Xanthine/xanthine oxidase (50mU/ml), H_2_O_2_=Hydrogen peroxide 200µM), 4D-after 4 days from start of experiment, 9D-after 9 days from start of experiment, 5DR-After 5 days of REM sleep recovery i.e. rats were allowed to sleep in cages for 5 days]

### Sleep loss increased the susceptibility of hepatocytes for oxidative stress

REM sleep loss not only generated ROS in the hepatocytes of rats but also made them more susceptible to oxidative stresses. We report here the increased production of ROS for REMSD group hepatocytes after day 4 (135.50±13.5%) and day 9 (257.1±15.75%) compared to CC groups, when hepatocytes were exposed to 200µM-H_2_O_2_ for 60 minutes (Fig.2b, ANOVA F=216.67, df=11, p<0.001 & Fig.2c, ANOVA F=442.44, df=15, p<0.001). The levels of ROS became normal when similar number of hepatocytes were treated with 500U/ml catalase (Fig. 2b &c). Similarly, when hepatocytes from CC, LPC and REMSD groups after day 4, 9 and after 5DR were treated with 50 mU/ml-X/XO, different levels of ROS were induced in them. The ROS levels observed after 4 day REMSD rat hepatocytes were 75.1±6.5% while after 9 day it became 130.5±5.5% of the control (Fig 2b and C). The treatment with 50 mU/ml-X/XO+50µM-Oxypurinol treatment reduced the ROS levels comparable to CC and LPD group of rats (Fig.2b&c). The ROS levels in hepatocytes collected from 5DR rats responded similar to CC and LPC group of rats (Fig. 2c).

### Expression of Nitic oxide synthase gene and protein circulation due to REM sleep deprivation

The iNOS gene expression got affected in hepatocytes of REMSD rats. We found 0.6±0.08 fold increased expression of iNOS gene on day 4 compared to CC control, while expression on day 9 increased up to 1.5±0.15 fold by 9^th^ day of REMSD. After 5days of recovery sleep, the expression pattern improved towards normal level (0.71±0.05 fold) compared to control (Fig.3a, ANOVA F=357.99, df=6, p<0.001). Further, we measured iNOS protein levels in hepatocytes using western blot under similar conditions. Our results shows that iNOS protein level increased 95.1±1.5% on day 4^th^, which subsequently increased to 162.5±1.25% after 9 days of REMSD compared to CC group of rats (Fig.3 b & c, ANOVA F=393.94, df=4, p<0.001). The levels of iNOS protein returned to normal levels after 5 days of recovery sleep (42.5±1.1%), compared to controls (Fig.3 b & c).

### Nitic oxide production in hepatocytes after REMSD

As production of iNOS protein increased in the hepatocytes due to REM sleep loss, we collected the hepatocytes from different groups of rats and measured NO levels in them. Our results suggest that, NO levels were significantly increased in hepatocytes of the REMSD group rats on day 4 (285±5%) and day 9 (675±11%) than that of cage control group. Further, 5 days of sleep recovery showed redressal effects in terms of NO production but overall NO level was still found higher (135±12%) than control (Fig. 4, ANOVA F=170.48, df=11, p<0.001).

**Figure 3:**
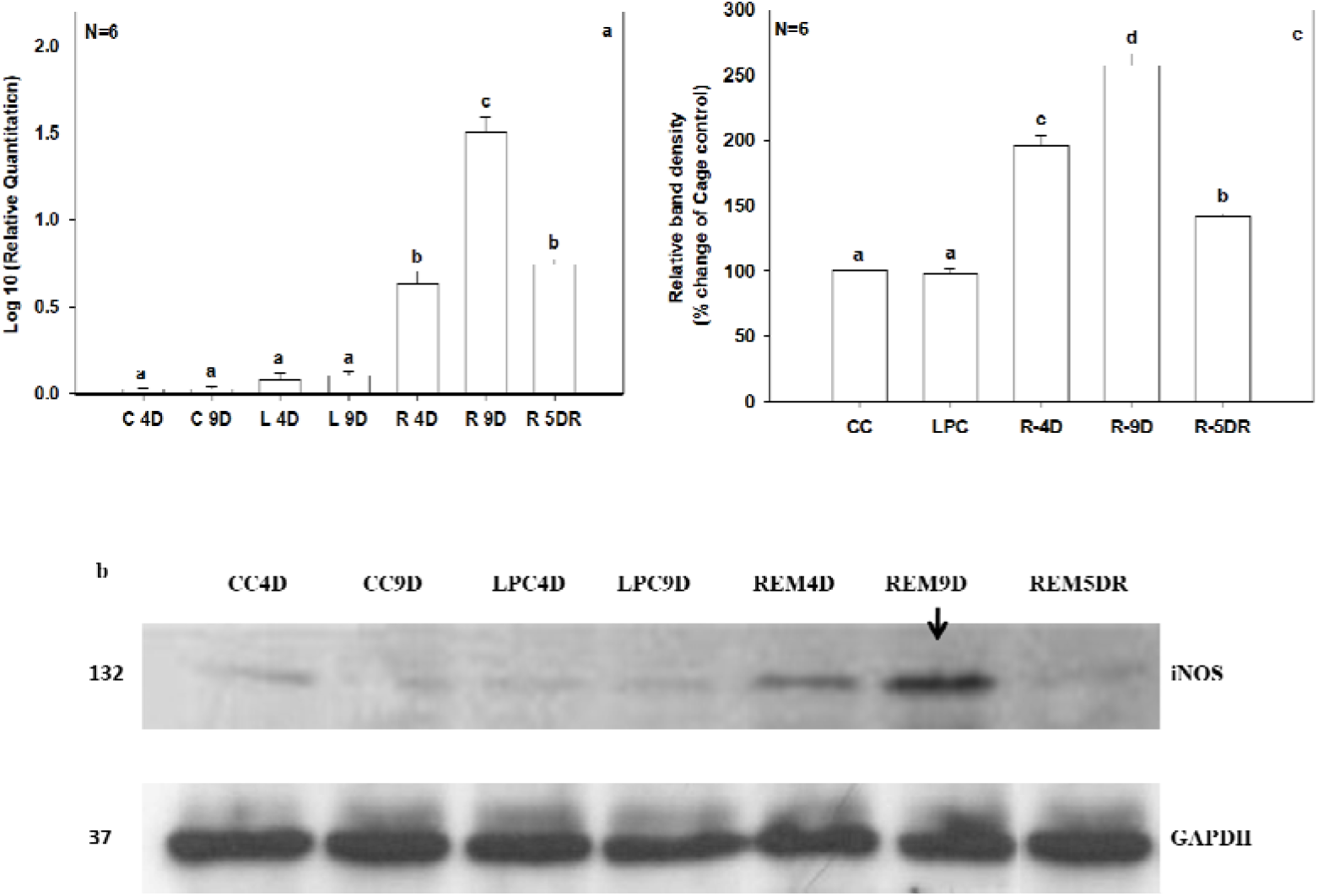
Measurement of iNOS gene and protein levels in response to REMSD in hepatocytes. **(a),** The graph shows log fold change in expression pattern of iNOS gene, cage control samples were taken as calibrator while GAPDH was endogenous control for respective genes. (b), Analysis of hepatocytes iNOS protein using WB from CC, LPC and REM sleep deprived and recovery group rats. (c), Densitometric analysis of WB samples for different treatment groups using Image-J software, NIH. Data were represented as relative band density reflected as percentage change of control. Refer figure 2 for statistical and legend details as P value < 0.05 were considered as statistically significant.

**Figure 4:**
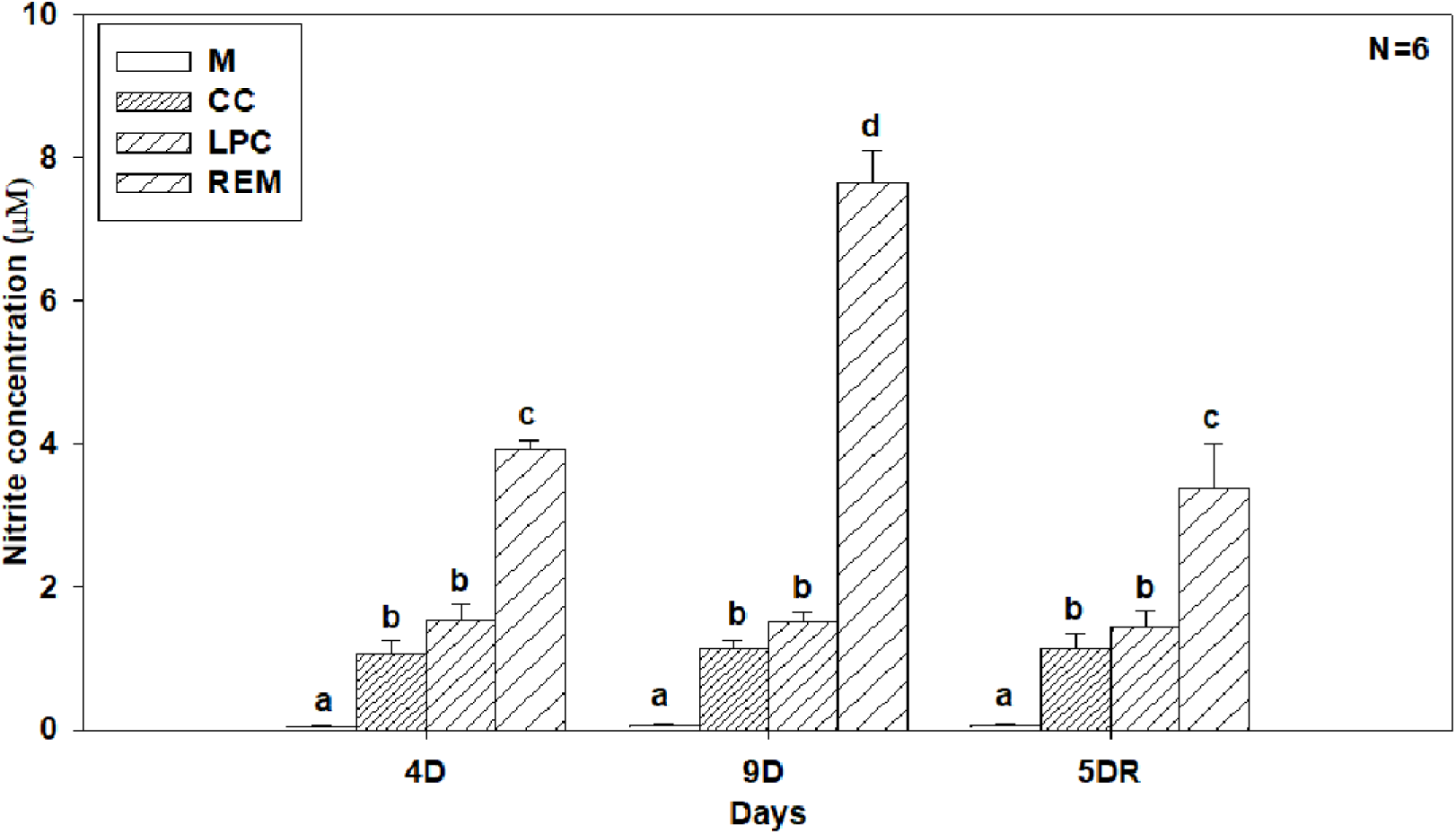
Measurement of NO level in hepatocytes in response to REM sleep loss and sleep recovery. Rats were REM sleep deprived of 4 days and 9 days before recovery of 5 days. The hepatocytes were collected from different time points and cultured for nitrite measurement. The X-axis represents the days of REM sleep deprivation and recovery and Y-axis represents the nitrite concentration (µM) in response to REM sleep deprivation. The Kit (Griess reagent kit, Sigma) was used to measure out the levels of nitrite from cultured hepatocytes. [M=Media only, CC=cage control, LPC=large platform control, REM=rapid eye movement), refer figure 2 for statistical details and other legend abbreviation].

## Discussions

We subjected the rats to REMSD using flower-pot method and created the LPC groups and CC groups under similar conditions for comparison. We quantitated the ROS level in the hepatocytes of all the groups by measuring DCF florescence using FACS. We collected the hepatocytes and standardized for induction and suppression of ROS by using standard enzymes like catalase or oxypurinol. Lastly, we measured the expression levels of iNOS gene and the protein as well as NO level in hepatocytes after REMSD. Our findings suggest that REMSD can increase ROS production in the hepatocytes of rats which ultimately can induce apoptotic cell death [34]. Treatment of hepatocytes with H_2_O_2_ and X/XO selectively induced production of ROS inside the hepatocytes which could be quenched by catalase or oxypurinol treatment. This gave us a tool to expose the hepatocytes from CC, LPC and REMSD groups and investigate how REMSD will affect their susceptibility to future oxidative stresses.

The oxidative stress created by treatments of H_2_O_2_ and X/XO induced the ROS levels more in the REMSD group of rats compared to CC and LPC group of rats indicating the REM sleep loss affects stress induced potentiation of hepatocytes. While, the hepatocytes from 5 days sleep recovery showed normal levels comparable to controls. This can be explained as follows, REMSD might have weakened out the natural cell response by creating more oxidative stress. The stress response got alleviated to certain extent when rats were allowed to have recovery sleep. Other studies also indicate the increased oxidative stress in brain cells of rats after REMSD [18,35], reviewed in for animal models [36]. Conditions like obstructive sleep apnea in humans and rodents were also found to be positively correlated with increased ROS in the system [37,38], while Parkinson disease associated with REM sleep loss were also found to contribute to the colon oxidative stress and mitochondrial function [39]. Sleep deprivation was further found to be associated with anxiety like behavior in rats which can be reduced or prevented by administration of melatonin, possibly by reducing the oxidative stress and maintain the balance between GABAergic and glutamatergic transmission [40]. Administrations of melatonin were found to reduce oxidative stress in rats by blocking the down-regulation of γ-aminobutyric acid A-alpha-2 receptors. These studies suggest that sleep deprivation is somehow directly or indirectly associated with the ROS levels in different tissues studied so far. In our previous study, we had also reported that REMSD can create inflammation in liver by increasing the levels of cytokines such as Interleukin-6 (IL-6), IL-12, IL-1β [13] indicating that cytokines might be involved in induction of iNOS genes. Results published earlier also suggest that cytokines like tumor necrosis factor alpha (TNF-α), IL-1β, and Interferon gamma (INF-γ) synergistically activate iNOS gene expression in the liver [35].

Lastly, our results suggest that hepatocytes form REMSD group of rats showed increased NO production which got reduced after sleep recovery. Previous studies claim that NO exerts a protective effect both *in vivo* and *in vitro* by blocking TNF-α induced apoptosis and hepatotoxicity, by inhibiting the caspase-3-like protease activity [41]. Thus, our unpublished data showing that REMSD can cause apoptosis like situation in hepatocytes and increased levels of NO in response to REMSD suggest that excess of NO were being produced due to sleep loss to counteract the oxidative stress generated due to sleep loss and protect the hepatocytes from apoptosis like processes [34]. Our study is quite unique in testing the oxidative response as a function of REMSD with optimized enzyme treatment protocols in rats which suggest that REM sleep deprivation can induce ROS in hepatocytes and similarly makes them more vulnerable to oxidative stress. This study further supports the idea of REM sleep demonstrating the functional role of maintenance and enhancing survival of individuals.

## Conclusions

Our results demonstrate that the REM sleep is very important for individual well-being and its loss can induce the hepatocytes for ROS production and further make them more susceptible to oxidative stress and finally can cause apoptotic cell death [34]. Apart from this, REMSD also can induce the iNOS circulation in the system inducing NO production, which might be directly or indirectly responsible for the wellbeing of hepatocytes.

## Abbreviations

ROS: reactive oxygen species
REM: rapid eye movement
NREM: non rapid eye movement
REMSD: rapid eye movement sleep deprivation
CC: cage control
LPC: large platform control
EDTA: Ethylenediaminetetraacetic acid
RIPA: Radio immunoprecipitation assay
FITC: Fluorescein isothiocyanate
NO: Nitric Oxide

## Conflict of Interests

There is no conflict of interest among authors.

## References

[1] Cirelli C, Tononi G. Is sleep essential? PLoS Biol 2008;6:1605–11. doi:10.1371/journal.pbio.0060216.

[2] Reutrakul S, Van Cauter E. Sleep influences on obesity, insulin resistance, and risk of type 2 diabetes. Metabolism 2018;84:56–66. doi:10.1016/j.metabol.2018.02.010.

[3] Tobaldini E, Costantino G, Solbiati M, Cogliati C, Kara T, Nobili L, et al. Sleep, sleep deprivation, autonomic nervous system and cardiovascular diseases. Neurosci Biobehav Rev 2017;74:321–9. doi:10.1016/j.neubiorev.2016.07.004.

[4] Yang H, Haack M, Gautam S, Meier-Ewert HK, Mullington JM. Repetitive exposure to shortened sleep leads to blunted sleep-associated blood pressure dipping. J Hypertens 2017;35:1187–94. doi:10.1097/HJH.0000000000001284.

[5] Spiegel K, Leproult R, Van Cauter E. Impact of sleep debt on metabolic and endocrine function. Lancet 1999;354:1435–9. doi:10.1016/S0140-6736(99)01376-8.

[6] Graves L, Heller E, Pack A, Abel T. Sleep deprivation selectively impairs memory consolidation for contextual fear conditioning. Learn Mem 2003;10:168–176. doi:10.1101/lm.48803.LEARNING.

[7] Kumar T, Jha SK. Sleep deprivation Impairs Consolidation of Cued Fear Memory in Rats. PLoS One 2012;7. doi:10.1371/journal.pone.0047042.

[8] Youngblood BD, Zhou J, Smagin GN, Ryan DH, Harris RBS. Sleep deprivation by the “flower pot” technique and spatial reference memory. Physiol Behav 1997;61:249–56. doi:10.1016/S0031-9384(96)00363-0.

[9] Mallick BN, Singh S, Pal D. Role of alpha and beta adrenoceptors in locus coeruleus stimulation-induced reduction in rapid eye movement sleep in freely moving rats. Behav Brain Res 2005;158:9–21. doi:10.1016/j.bbr.2004.08.004.

[10] Gómez-González B, Hurtado-Alvarado G, Esqueda-León E, Santana-Miranda R, Rojas-Zamorano JÁ, Velázquez-Moctezuma J. REM sleep loss and recovery regulates blood-brain barrier function. Curr Neurovasc Res 2013;10:197–207. doi:10.2174/15672026113109990002.

[11] Christos GA. Is Alzheimer’s disease related to a deficit or malfunction of rapid eye movement (REM) sleep? Med Hypotheses 1993;41:435–9. doi:10.1016/0306-9877(93)90121-6.

[12] Baumann C, Ferini-Strambi L, Waldvogel D, Werth E, Bassetti CL. Parkinsonism with excessive daytime sleepiness--a narcolepsy-like disorder? J Neurol 2005;252:139–45. doi:10.1007/s00415-005-0614-5.

[13] Maquet P. Functional neuroimaging of normal human sleep by positron emission tomography. J Sleep Res 2000;9:207–31. doi:jsr214 [pii].

[14] Everson C a, Smith CB, Sokoloff L. Effects of prolonged sleep deprivation on local rates of cerebral energy metabolism in freely moving rats. J Neurosci 1994;14:6769–78.

[15] Mejri MA, Yousfi N, Hammouda O, Tayech A, Ben Rayana MC, Driss T, et al. One night of partial sleep deprivation increased biomarkers of muscle and cardiac injuries during acute intermittent exercise. J Sports Med Phys Fitness 2017;57:643–51. doi:10.23736/S0022-4707.16.06159-4.

[16] Silva RH, Abílio VC, Takatsu AL, Kameda SR, Grassl C, Chehin AB, et al. Role of hippocampal oxidative stress in memory deficits induced by sleep deprivation in mice. Neuropharmacology 2004;46:895–903. doi:10.1016/j.neuropharm.2003.11.032.

[17] Everson CA, Laatsch CD, Hogg N. Antioxidant defense responses to sleep loss and sleep recovery. Am J Physiol Regul Integr Comp Physiol 2005;288:R374–383. doi:10.1152/ajpregu.00565.2004.

[18] Gopalakrishnan A, Ji LL, Cirelli C. Sleep deprivation and cellular responses to oxidative stress. Sleep 2004;27:27–35. doi:10.1093/sleep/27.1.27.

[19] D’Almeida V, Lobo LL, Hipólide DC, de Oliveira a C, Nobrega JN, Tufik S. Sleep deprivation induces brain region-specific decreases in glutathione levels. Neuroreport 1998;9:2853–6. doi:10.1097/00001756-199808240-00031.

[20] Ramanathan L, Gulyani S, Nienhuis R, Siegel JM. Sleep deprivation decreases superoxide dismutase activity in rat hippocampus and brainstem. Neuroreport 2002;13:1387–90. doi:10.1097/00001756-200208070-00007.

[21] D’Almeida V, Hipolide DC, Azzalis LA, Lobo LL, Junqueira VB, Tufik S. Absence of oxidative stress following paradoxical sleep deprivation in rats. Neurosci Lett 1997;235:25–8.

[22] Brown MK, Naidoo N. The UPR and the anti-oxidant response: Relevance to sleep and sleep loss. Mol Neurobiol 2010;42:103–13. doi:10.1007/s12035-010-8114-8.

[23] Chang HM, Mai F Der, Chen BJ, Wu UI, Huang YL, Lan CT, et al. Sleep deprivation predisposes liver to oxidative stress and phospholipid damage: A quantitative molecular imaging study. J Anat 2008;212:295–305. doi:10.1111/j.1469-7580.2008.00860.x.

[24] Rui L. Energy metabolism in the liver. Compr Physiol 2014;4:177–97. doi:10.1002/cphy.c130024.

[25] Ménochet K, Kenworthy KE, Houston JB, Galetin A. Simultaneous assessment of uptake and metabolism in rat hepatocytes: a comprehensive mechanistic model. J Pharmacol Exp Ther 2012;341:2–15. doi:10.1124/jpet.111.187112.

[26] Rogiers V, Vandenberghe Y, Callaerts A, Verleye G, Cornet M, Mertens K, et al. Phase I and phase II xenobiotic biotransformation in cultures and co-cultures of adult rat hepatocytes. Biochem Pharmacol 1990;40:1701–6. doi:10.1016/0006-2952(90)90345-L.

[27] Hutson SM, Stinson-Fisher C, Shiman R, Jefferson LS. Regulation of albumin synthesis by hormones and amino acids in primary cultures of rat hepatocytes. Am J Physiol Metab 1987;252:E291–8. doi:10.1152/ajpendo.1987.252.3.E291.

[28] Pandey AK, Kar SK. REM sleep deprivation of rats induces acute phase response in liver. Biochem Biophys Res Commun 2011;410. doi:10.1016/j.bbrc.2011.05.123.

[29] Hicks RA, Okuda A, Thomsen D. Depriving rats of REM sleep: the identification of a methodological problem. Am J Psychol 1977;90:95–102. doi:10.2307/1421644.

[30] van Hulzen ZJM, Coenen AML. Paradoxical sleep deprivation and locomotor activity in rats. Physiol Behav 1981;27:741–4. doi:10.1016/0031-9384(81)90250-X.

[31] Liu XL, Li LJ, Chen Z. Isolation and primary culture of rat hepatocytes. Hepatobiliary Pancreat Dis Int 2002;1:77–79. doi:10.1007/BF00303186.

[32] Bai X chun, Lu D, Liu A ling, Zhang Z ming, Li X mei, Zou Z peng, et al. Reactive Oxygen Species Stimulates Receptor Activator of NF-{kappa}B Ligand Expression in Osteoblast. J Biol Chem 2005;280:17497–506. doi:10.1074/jbc.M409332200.

[33] Massaad C a, Portier BP, Taglialatela G. Inhibition of transcription factor activity by nuclear compartment-associated Bcl-2. J Biol Chem 2004;279:54470–8. doi:10.1074/jbc.M407659200.

[34] Atul Pandey; Devesh Kumar; Gopesh Ray; Santosh Kar. Rapid eye movement sleep deprivation causes apoptotic cell-death of the hepatocytes in rat. Biorxiv 2018. doi:https://doi.org/10.1101/375717.

[35] Mathangi DC, Shyamala R, Subhashini AS. Effect of REM sleep deprivation on the antioxidant status in the brain of Wistar rats. Ann Neurosci 2012;19:161–4. doi:10.5214/ans.0972.7531.190405.

[36] Villafuerte G, Miguel-Puga A, Rodriguez EM, Machado S, Manjarrez E, Arias-Carrion O. Sleep deprivation and oxidative stress in animal models: a systematic review. Oxid Med Cell Longev 2015;2015:234952. doi:10.1155/2015/234952.

[37] Eisele H-J, Markart P, Schulz R. Obstructive Sleep Apnea, Oxidative Stress, and Cardiovascular Disease: Evidence from Human Studies. Oxid Med Cell Longev 2015;2015:1–9. doi:10.1155/2015/608438.

[38] Wang Y, Zhang SX, Gozal D. Reactive oxygen species and the brain in sleep apnea. Respir Physiol Neurobiol 2010;174:307–16. doi:10.1016/j.resp.2010.09.001.

[39] Morén C, González-Casacuberta, Navarro-Otano J, Juárez-Flores D, Vilas D, Garrabou G, et al. Colonic Oxidative and Mitochondrial Function in Parkinson’s Disease and Idiopathic REM Sleep Behavior Disorder. Parkinsons Dis 2017;2017. doi:10.1155/2017/9816095.

[40] Zhang L, Guo HL, Zhang HQ, Xu TQ, He B, Wang ZH, et al. Melatonin prevents sleep deprivation-associated anxiety-like behavior in rats: Role of oxidative stress and balance between gabaergic and glutamatergic transmission. Am J Transl Res 2017;9:2231–42.

[41] Taylor BS, Alarcon LH, Billiar TR. Inducible nitric oxide synthase in the liver: Regulation and function. Biochem Engl Tr 1998;63:766–81.

